# A Human Tissue-Specific Transcriptomic Analysis Reveals that Ageing Hinders Cancer and Boosts Cellular Senescence

**DOI:** 10.1101/595041

**Authors:** Kasit Chatsirisupachai, Daniel Palmer, Susana Ferreira, João Pedro de Magalhães

**Author notes:** **Corresponding author** João Pedro de Magalhães, Integrative Genomics of Ageing Group, Institute of Ageing and Chronic Disease, University of Liverpool, William Duncan Building, Room 281, 6 West Derby Street, Liverpool L7 8TX, United Kingdom.

## Abstract

Ageing is the biggest risk factor for cancer, but the mechanisms linking these two processes remain unclear. We compared genes differentially expressed with age and genes differentially expressed in cancer among nine human tissues. In most tissues, ageing and cancer gene expression surprisingly changed in the opposite direction. These overlapping gene sets were related to several processes, mainly cell cycle and the immune system. Moreover, cellular senescence signatures derived from a meta-analysis changed in the same direction as ageing and in the opposite direction of cancer signatures. Therefore, transcriptomic changes in ageing and cellular senescence might relate to a decrease in cell proliferation, while cancer transcriptomic changes shift towards an increase in cell division. Our results highlight the complex relationship between ageing, cancer and cellular senescence and suggest that in most human tissues ageing processes and senescence act in tandem while being detrimental to cancer. Our work challenges the traditional view concerning the relationship between cancer and ageing and suggests that ageing processes may hinder cancer development.

## MAIN TEXT

Ageing is the biggest risk factor for cancer (de Magalhaes 2013). However, the biological mechanisms behind this link are still unclear. Gene expression analyses have been used to study cancer (Cieslik & Chinnaiyan 2018) and ageing (de Magalhaes *et al.* 2009; Yang *et al.* 2015), but only a few studies have investigated the relationship between gene expression changes in these two processes (Aramillo Irizar *et al.* 2018). In particular, comparisons between human tissue-specific genes differentially expressed with age (age-DEGs) and genes differentially expressed in cancer (cancer-DEGs) are lacking. Cellular senescence is a state of irreversible cell cycle arrest and has been regarded as an anti-tumour mechanism (Campisi 2013). Many studies have attempted to identify gene expression signatures of senescence (Kim *et al.* 2013), but comparisons between cellular senescence signatures and age-DEGs and cancer-DEGs are also lacking.

Recently, the Genotype-Tissue Expression (GTEx) Project has profiled gene expression from non-cancerous tissues of nearly 1,000 individuals over 53 sampled sites (Consortium 2015). The Cancer Genome Atlas (TCGA) has sequenced tumour samples from more than 10,000 patients covering 33 cancer types (Ding *et al.* 2018). Here, we investigated the relationship between transcriptomic changes in human tissue ageing and cancer by analyzing gene expression data from GTEx and TCGA. In addition, we conducted a meta-analysis using publicly available datasets to identify cellular senescence signature genes and compared them with age-DEGs and cancer-DEGs.

We first identified age-DEGs in nine tissues from GTEx (Supporting Information, Table S1). The numbers of significant age-DEGs (FDR < 0.05 and absolute fold change > 1.5; moderated t-test) varied between different tissues (Figure 1a, Data S1). We identified cancer-DEGs by analyzing nine TCGA datasets for which the tissues of origin were matched to the GTEx tissues used in this study (Supporting Information, Table S1). The numbers of cancer-DEGs (FDR < 0.01 and absolute fold change > 2; moderated t-test) are shown in Figure 1b (Data S2).

**Figure 1.**
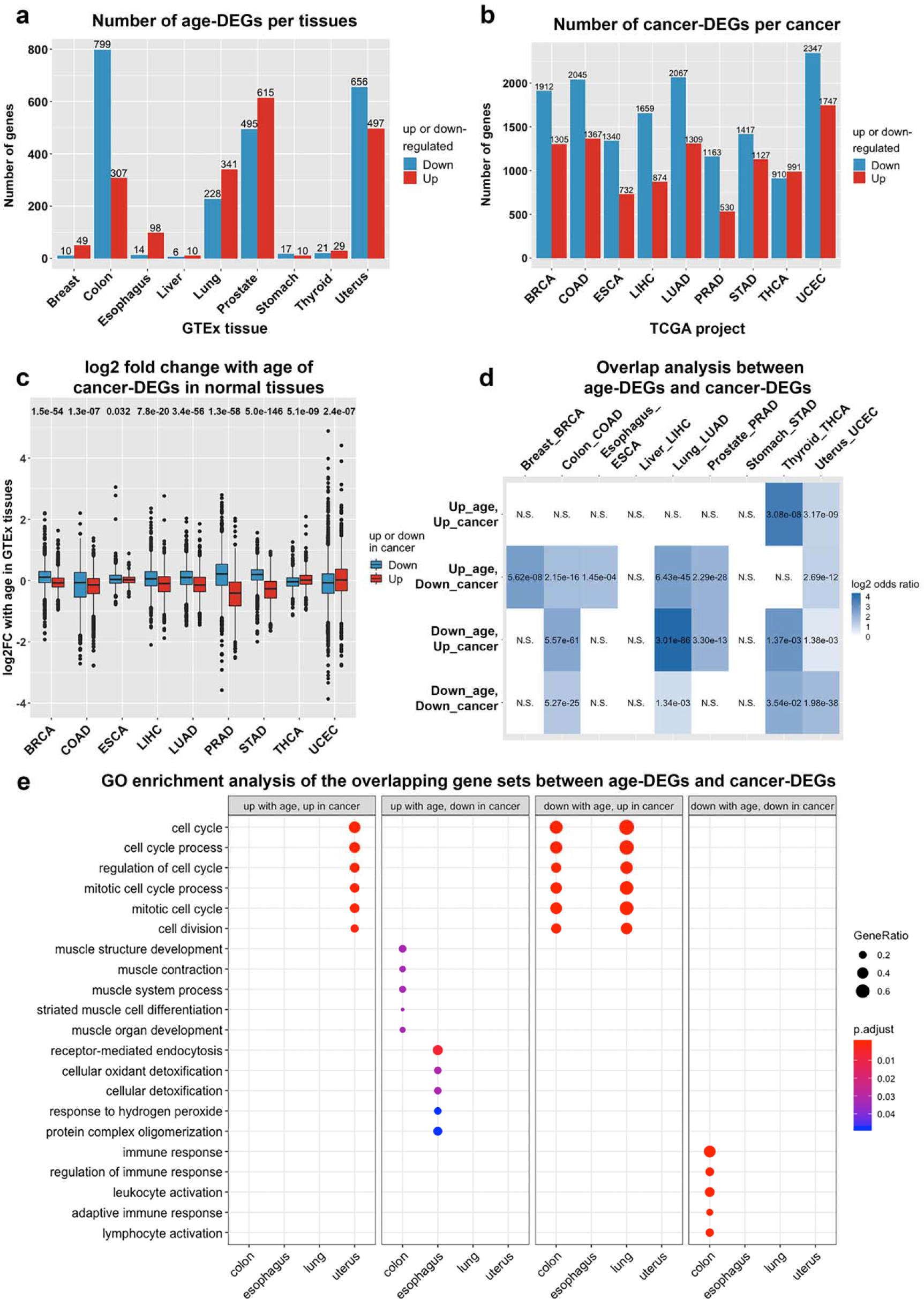
The relationship between age-DEGs and cancer-DEGs. (a) Number of age-DEGs. (b) Number of cancer-DEGs. (c) Fold change with age in GTEx data of cancer-DEGs. Numbers indicate *P*-values (Mann-Whitney *U* test). (d) Overlap between age-DEGs and cancer-DEGs. Numbers represent FDR (Fisher’s exact test, Benjamini-Hochberg correction). N.S. denotes non-significant overlap. Colours of the heatmap correspond to odds ratio. (e) GO enrichment analysis of significant overlap gene sets. The plot shows examples of significant enriched terms (FDR < 0.1).

After obtaining a list of cancer-DEGs, we examined the fold-change with age of the cancer-DEGs. We observed a significantly higher fold-change with age in genes down-regulated with cancer when compared to genes up-regulated with cancer for most cancer types, with the opposite being observed in two tissues: THCA-thyroid and UCEC-uterus (*P*-value < 0.05; Mann-Whitney *U* test) (Figure 1c). This result may partly explain the reduction in cancer incidence and mortality in the oldest old (Aramillo Irizar *et al.* 2018). We next overlapped age-DEGs and cancer-DEGs for each tissue. Overall, genes changed in the opposite directions between ageing and cancer significantly overlapped more often than genes changed in the same direction (Figure 1d). There was no significant overlap in liver and stomach, which might be explained by the low number of age-DEGs. In uterus, however, the overlap was significant in all cases.

We performed GO enrichment analysis and found that 6 out of 20 significantly overlapping sets were enriched in GO terms (Figure 1e, Data S3). Genes down-regulated with age – up-regulated in cancer in colon and lung were related to cell cycle. Cell cycle terms were also enriched in genes up-regulated with age – up-regulated in cancer in uterus. Uncontrolled cell proliferation in the ageing uterus often leads to endothelial hyperplasia and could lead to endometrial cancer (Damle *et al.* 2013). Immune-related terms were enriched in genes down-regulated with age – down-regulated in cancer in the colon. The immune system plays an important role in preventing cancer through immunosurveillance (Ribatti 2017). Therefore, compromised immune function with age would be likely to provide the immunosuppressive microenvironment, which allows cancer cells to evade immunosurveillance. Overall, these results highlight the tissue specificity of processes altered during ageing and cancer.

Cancer driver mutations accumulate with age and may initiate tumourigenesis (Blokzijl *et al.* 2016). While mutations are the main driver of cancer, our results suggest that ageing processes may actually hinder cancer development. Ageing and other age-related diseases, such as neurodegenerative diseases, are involved in the loss of function and degeneration of the cells, yet cancer cells rely on the gain-of-function phenotypes including an increase in proliferation. Our results fit well with this notion and highlight that the relationship between cancer and ageing is far from straightforward.

We next performed a meta-analysis using 20 microarray datasets from Gene Expression Omnibus (GEO) and identified 526 overexpressed and 734 underexpressed genes (Supporting Information, Table S2, Data S4). GO enrichment analysis indicated that underexpressed signatures were related to cell cycle, while overexpressed signatures were linked to immune response processes (Figure S1, Data S5). KEGG pathway enrichment revealed that underexpressed senescence signature genes were related to cell cycle and DNA repair. Overexpressed senescence signature genes were associated with lysosome and p53 signaling pathway, the well-established senescence pathway (Figure S2, Data S5). Thus, we have identified cellular senescence signature genes that are consistently over- or underexpressed across datasets, and showed that these genes were associated with different senescence-related pathways.

Underexpressed senescence signatures overlapped with genes down-regulated with age in the colon and lung, while overexpressed senescence signatures overlapped with genes up-regulated with age in colon, lung, prostate, and thyroid (Figure 2a-b). These results provide evidence that senescent cells accumulate with age in human tissues. Only the uterus displayed opposite results, in line with the up-regulation in the ageing uterus of cell cycle-related genes (Figure 2a-b). For cancer genes, as expected, an opposite trend between cancer-DEGs and cellular senescence signatures emerged (Figure 2c-d). This highlights the anti-cancer role of senescence. Only in THCA-thyroid were the genes up-regulated in cancer significantly overlapped with overexpressed senescence genes.

**Figure 2.**
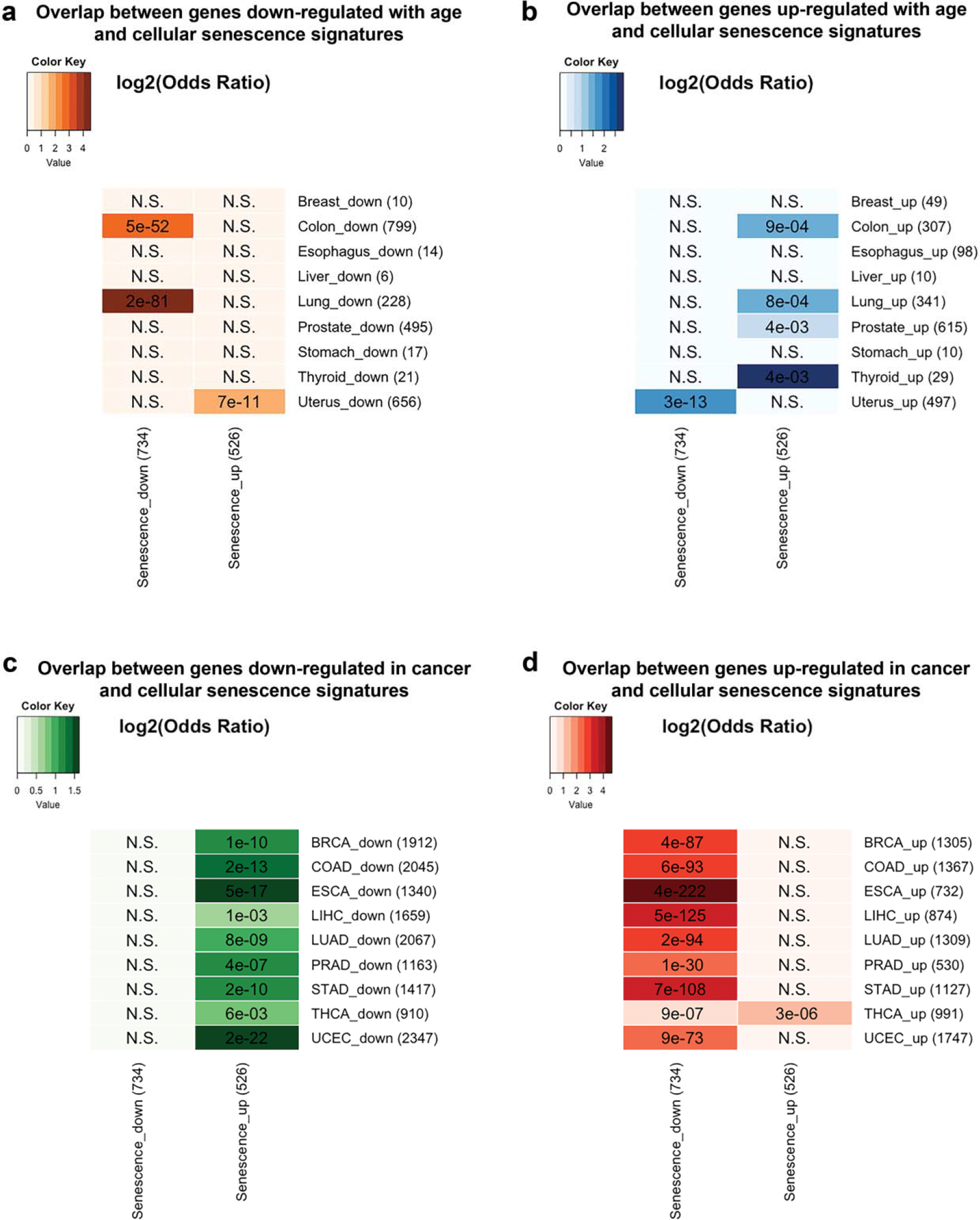
Overlap between cellular senescence signature genes and genes (a) down-regulated with age, (b) up-regulated with age, (c) down-regulated in cancer, and (d) up-regulated in cancer. Numbers represent FDR (Fisher’s exact test, Benjamini-Hochberg correction). N.S. denotes non-significant overlap. Colours correspond to log2 odds ratio.

In conclusion, our study shows that transcriptomic changes with age tend to be associated with cellular senescence signatures, but opposite to cancer in most tissues, with the exception of the uterus. The overlapping genes between cancer and ageing were linked to different processes mainly related to cell cycle. The gene lists provided by our study could also serve as a resource for future experimental studies. This study provides novel insights into the complex relationship between transcriptomic changes in human ageing, cancer, and cellular senescence. Our work challenges the traditional view concerning the relationship between cancer and ageing.

## Acknowledgements

Work in our lab is supported by the Wellcome Trust (208375/Z/17/Z), the Leverhulme Trust (RPG-2016-015) and the Biotechnology and Biological Sciences Research Council (BB/R014949/1). KC is supported by a Mahidol-Liverpool PhD scholarship from Mahidol University, Thailand, and the University of Liverpool, UK.

## References

Aramillo IrizarP, SchaubleS, EsserD, GrothM, FrahmC, PriebeS, … Kaleta C (2018). Transcriptomic alterations during ageing reflect the shift from cancer to degenerative diseases in the elderly. Nat Commun. 9, 327.

BlokzijlF, de LigtJ, JagerM, SasselliV, RoerinkS, SasakiN, … van BoxtelR (2016). Tissue-specific mutation accumulation in human adult stem cells during life. Nature. 538, 260–264.

Campisi J (2013). Aging, cellular senescence, and cancer. Annu Rev Physiol. 75, 685–705.

CieslikM, Chinnaiyan AM (2018). Cancer transcriptome profiling at the juncture of clinical translation. Nat Rev Genet. 19, 93–109.

Consortium GT (2015). Human genomics. The Genotype-Tissue Expression (GTEx) pilot analysis: multitissue gene regulation in humans. Science. 348, 648–660.

DamleRP, DravidNV, SuryawanshiKH, GadreAS, BagalePS, Ahire N (2013). Clinicopathological Spectrum of Endometrial Changes in Peri-menopausal and Post menopausal Abnormal Uterine Bleeding: A 2 Years Study. J Clin Diagn Res. 7, 2774–2776.

de Magalhaes JP (2013). How ageing processes influence cancer. Nat Rev Cancer. 13, 357–365.

de MagalhaesJP, CuradoJ, Church GM (2009). Meta-analysis of age-related gene expression profiles identifies common signatures of aging. Bioinformatics. 25, 875–881.

DingL, BaileyMH, Porta-PardoE, ThorssonV, ColapricoA, BertrandD, … Cancer Genome Atlas Research N (2018). Perspective on Oncogenic Processes at the End of the Beginning of Cancer Genomics. Cell. 173, 305–320 e310.

KimYM, ByunHO, JeeBA, ChoH, SeoYH, KimYS, … Yoon G (2013). Implications of time-series gene expression profiles of replicative senescence. Aging Cell. 12, 622–634.

Ribatti D (2017). The concept of immune surveillance against tumors. The first theories. Oncotarget. 8, 7175–7180.

YangJ, HuangT, PetraliaF, LongQ, ZhangB, ArgmannC, … Tu Z (2015). Synchronized age-related gene expression changes across multiple tissues in human and the link to complex diseases. Sci Rep. 5, 15145.

