## Supplementary material for "A Human Tissue-Specific Transcriptomic Analysis Reveals that Ageing Hinders Cancer and Boosts Cellular Senescence"

### Supporting Information

### Methods

#### Tissue-specific genes differentially expressed with age (age-DEGs) from GTEx

The RNA-Seq based gene expression data of non-cancerous tissues (v7, January 2015 release) were download from the GTEx portal (https://gtexportal.org) (Consortium 2015). Out of 30 tissues provided by GTEx, four tissues (bladder, cervix uteri, fallopian tube, and kidney) with low samples (11, 11, 7, and 45, respectively) were excluded from the analysis. For each tissue, we identified differentially expressed genes with age using the following linear regression model:

$$Y_{ij} = \alpha Age_i + \beta Sex_i + \gamma Death_i$$

Where $Y_{ij}$ is the expression level of gene $j$ in sample $i$, $Age_i$ denotes the age of sample $i$. $Sex_i$ denotes the sex of sample $I$, $Death_i$ denotes the death classification of sample $i$ based on the 4-point Hardy scale (Ferreira *et al.* 2018). It should be noted that dataset downloaded from GTEx portal did not provide the actual age of each sample, the age ranges, i.e. 20-29, 30-39, 40-49, 50-59, 60-69 and 70-79, were provided instead. We then approximated the age of each sample to 25, 35, 45, 55, 65 and 75, respectively. Because our main aim was to compare the age-DEGs with cancer-DEGs from TCGA, and there was a large difference between the number of the total genes in GTEx (56,202) and TCGA (20,532), we focus only on protein coding genes. Protein-coding genes were identified using the R package *biomaRt* (version 2.36.1) (Durinck *et al.* 2009), based on Ensembl release 92 (April 2018). After removing the non-coding genes, there were 18,851 protein-coding genes. To remove the low expressed genes, genes with expression less than 1 count per million (cpm) in more than 30 percent of samples were excluded. Raw read counts were normalized using TMM normalization and were voom transformed to remove heteroscedasticity from the count data. The linear model for each gene was generated by using the *limma* package in R (version 3.36.5). Genes were considered to be significant differentially expressed genes with age if the empirical Bayes moderated t-statistics and their associated adjust $P$-value (Benjamini-Hochberg method) < 0.05 and absolute fold change across 50 years of age (from 25 to 75 years old) > 1.5. The genes in GTEx were in ensemble id, we then converted them into entrez id format using *biomaRt*. Lists of age-DEGs from all 26 tissues are provided in the Data S1.

We selected nine tissues for further analysis, including breast, colon, esophagus, liver, lung, prostate, stomach, thyroid and uterus. This selection was based on the criteria that 1) each of these tissues had a matched TCGA project from the same tissue of origin and 2) the number of solid normal samples in those TCGA projects were more than 10. Numbers of samples used in our study were summarized in Table S1. Most of the age-DEGs were tissue-specific, indicating the tissue specificity of age-DEGs, consistent with a previous report (Yang *et al.* 2015).

**Genes differentially expressed in cancer (cancer-DEGs) from nine TCGA projects**

The TCGA level 3 RNA-Seq based gene expression data for nine TCGA projects used in our analysis (BRCA (Cancer Genome Atlas 2012b), COAD (Cancer Genome Atlas 2012a), ESCA (Cancer Genome Atlas Research *et al.* 2017), LIHC (Cancer Genome Atlas Research Network. Electronic address & Cancer Genome Atlas Research 2017), LUAD (Cancer Genome Atlas Research 2014b), PRAD (Cancer Genome Atlas Research 2015), STAD (Cancer Genome Atlas Research 2014a), THCA (Cancer Genome Atlas Research 2014c), and UCEC (Cancer Genome Atlas Research *et al.* 2013)) were download from FireBrowse (http://firebrowse.org) in August 2018. RNA sequencing was performed on the Illumina Hi-Seq platform and processed using RNASeqV2 pipeline of TCGA, which provide the RSEM expected counts. Protein-coding genes were identified using the *biomaRt* R package (version 2.36.1). After removing non-coding genes, there were 18,583 protein-coding genes included in subsequent analysis. To remove the genes with low expression, genes with expression less than 1 count per million (cpm) in more than 30 percent of samples were excluded. The *limma* package in R was used to identify genes significantly differentially expressed between cancer and normal samples. The moderated t statistic *P*-value after correction by Benjamini-Hochberg method < 0.01 with an absolute fold-change > 2 were used to determine the statistical significance. Lists of cancer-DEGs from all 26 tissues were provided in the supplementary Data S2.

**Fold change with age in normal tissues of cancer-DEGs**

We examined the fold change with age of the cancer-DEGs. After obtaining the cancer-DEGs from TCGA, we identified the fold change with age in normal tissues in GTEx of these cancer-DEGs. Although not every cancer-DEG was found in the GTEx data from the corresponding tissue because lowly expressed genes had been filtered out before performing linear regression, the majority of them retained (Data S6). For each tissue, the fold change with age in normal tissue of up-regulated genes in cancer and down-regulated genes in cancer were compared using Mann-Whitney *U* test.

**Meta-analysis to identify cellular senescence signature genes**

To identify cellular senescence signature genes, the meta-analysis of 20 cellular senescence GEO datasets was performed. The datasets included in the analysis were shown in Table S2. We first separately identified differentially expressed genes in each dataset. The probe annotation for each dataset was slightly different, depended on the platform. In general, the steps are 1) Download the dataset from GEO, 2) Probe annotation to match probe ID with entrez ID using either the dataset's own GPL platform files, R packages (*AnnotationDbi* version 1.44.0 and *org.Hs.eg.db* version 3.7.0), or bioDBnet (https://biodbnet-abcc.ncifcrf.gov/db/db2db.php) (Mudunuri *et al.* 2009) 3) Remove probes which do not match to any entrez ID 4) Remove probes which match to multiple entrez ID 5) Average intensity of probes which match to the same entrez ID. The differential expression analysis was performed using limma. Genes with a *P*-value below 0.05 and absolute fold change > 1.5 were considered putatively differentially expressed genes. For the datasets with only one non-senescent sample and one senescent sample, only fold change cut-off was used to determine differentially expressed genes. There are two platforms (GPL2937 and GPL2947) in the dataset GSE3460, so we separately analysed each platform and treated them as two different datasets in the meta-analysis. After that, the results from all datasets were then combined using a binomial distribution and the false discovery rate set at $Q < 0.05$ using the same method as (de Magalhaes *et al.* 2009). In total, there were 1,259 senescence signature genes, with 526 overexpressed and 734 underexpressed. We found that 442 of 1,259 senescence signature genes were significantly overlapped with the signature of replicative senescence provided by Hernandez-Segura et al. (Hernandez-Segura *et al.* 2017) (*P*-value = 2.9e-161; Fisher's exact test), indicating the reliability of our data.

**Overlap analysis**

The overlap analyses were performed using the *GeneOverlap* package in R (version 1.16.0). In each experiment, background was set differently depending on where the genes come from. For the overlap between age-DEGs from GTEx and cancer-DEGs from TCGA, all protein coding genes in GTEx data (18,851 genes) were used as a background. For the overlap between age-DEGs and cellular senescence signature genes, all protein coding genes in combined datasets used for the meta-analysis (18,878 genes) were used as a background.

For the overlap between cancer-DEGs and cellular senescence signature genes, all protein coding genes in combined datasets used for the meta-analysis (18,878 genes). The overlap was considered significant if an adjusted *P*-value < 0.05 (Fisher's exact test followed by Benjamini-Hochberg correction).

**Functional enrichment analysis**

Gene ontology (GO) enrichment analysis was performed using the *clusterProfiler* package in R (version 3.8.1) (Yu *et al.* 2012). A GO term was considered to be an enriched term if an adjusted *P*-value < 0.1 (Benjamini-Hochberg correction). The union set of age-DEGs and cancer-DEGs in each of the four conditions for each tissue was employed as a background for GO enrichment test of the overlapping gene set in that tissue. For example, the union set of genes up-regulated with age in colon and genes up-regulated in COAD was used as a background for GO enrichment analysis of overlapping genes between genes up-regulated with age in colon and genes up-regulated in COAD. KEGG pathway enrichment analysis was performed on the underexpressed and overexpressed cellular senescence signatures. A pathway was enriched if an adjusted *P*-value < 0.05 (Benjamini-Hochberg correction). All protein coding genes in combined datasets used for the meta-analysis (18,878 genes) were employed as a background.

**List of Supplementary Tables, Supplementary Figures and Additional Data**

- Table S1 – Number of GTEx and TCGA samples used in the study
- Table S2 – GEO datasets for meta-analysis of cellular senescence signatures
- Figure S1 – GO enrichment analysis of cellular senescence signatures
- Figure S2 – KEGG pathway enrichment analysis of cellular senescence signatures
- Data S1 – List of age-DEGs in GTEx tissues
- Data S2 – List of cancer-DEGs in nine TCGA projects
- Data S3 – List of enriched terms from GO enrichment analysis of overlap genes between age-DEGs and cancer-DEGs
- Data S4 – List of cellular senescence signature genes
- Data S5 – GO and KEGG pathway enrichment analysis of cellular senescence signature genes
- Data S6 – Lists and fold change with age of cancer-DEGs presented in GTEx data

**Supplementary Table S1 – Number of GTEx and TCGA samples included in this study**

| **GTEx Tissues** | | **TCGA Projects** | | | |
|---|---|---|---|---|---|
| | | | | Samples | |
| Tissue Name | Cases | Disease Name | Project | Primary Solid Tumour | Solid Tissue Normal |
| Breast | 290 | Breast invasive carcinoma | BRCA | 1093 | 112 |
| Colon | 506 | Colon adenocarcinoma | COAD | 457 | 41 |
| Esophagus | 1018 | Esophageal carcinoma | ESCA | 184 | 11 |
| Liver | 174 | Liver hepatocellular carcinoma | LIHC | 371 | 50 |
| Lung | 425 | Lung adenocarcinoma | LUAD | 515 | 59 |
| Prostate | 152 | Prostate adenocarcinoma | PRAD | 497 | 52 |
| Stomach | 261 | Stomach adenocarcinoma | STAD | 415 | 35 |
| Thyroid | 444 | Thyroid carcinoma | THCA | 501 | 59 |
| Uterus | 110 | Uterine Corpus Endometrial Carcinoma | UCEC | 545 | 35 |

**Supplementary Table S2 – GEO datasets for meta-analysis of cellular senescence signatures**

| GEO | Platform ID | References | Tissue | Cell Type/ Cell Line | Proliferating Cells | Senescent Cells |
|---|---|---|---|---|---|---|
| GSE17077 | GPL1352 | (Gruber *et al.* 2010) | Annulus | - | 8 | 8 |
| GSE13330 | GPL570 | (Pazolli *et al.* 2009) | Foreskin | BJ Fibroblasts | 6 | 6 |
| GSE49860 | GPL5639 | (Imai *et al.* 2014) | - | Diploid Fibroblasts | 1 | 1 |
| GSE41714 | GPL10558 | (Kim *et al.* 2013) | Dermal | Diploid Fibroblasts | 2 (2 days) | 2 (30 and >30 days) |
| GSE37091 | GPL6480 | (Jong *et al.* 2013) | Umbilical vein | Endothelial cell (HUVECS) | 2 | 2 |
| GSE54095 | GPL6244 | (Guerrero *et al.* 2015) | Coronary Artery | Endothelial Cells | 4 | 4 |
| GSE3460 | GPL2937 | (Schwarze *et al.* 2002) | Prostate | Epithelial Cells | 2 | 2 |
| GSE3460 | GPL2947 | (Schwarze *et al.* 2002) | Prostate | Epithelial Cells | 1 | 1 |
| GSE3731 | GPL2990 | (Zhang *et al.* 2003) | Mammary | Epithelial Cells (HMEC) (48R and 184) | 8 | 8 |
| GSE36640 | GPL570 | (Shah *et al.* 2013) | - | Fibroblast/IMR90 | 5 | 5 |
| GSE19018 | GPL570 | - | - | Fibroblast/IMR90 | 3 | 3 |
| GSE28863 | GPL5175 | (Cao *et al.* 2011) | - | Fibroblasts | 12 | 12 |
| GSE3730 | GPL2990 | (Zhang *et al.* 2003) | - | Fibroblasts (WS1, WI38 and BJ) | 13 | 13 |
| GSE56530 | GPL570 | (Medeiros Tavares Marques *et al.* 2017) | Umbilical Cord Vein | Mesenchymal Stem Cells | 9 | 9 |
| GSE35957 | GPL570 | (Benisch *et al.* 2012) | Bone Marrow | Mesenchymal Stem Cells | 5 | 5 |
| GSE46019 | GPL6244 | (Frobel *et al.* 2014) | Bone Marrow | Mesenchymal Stromal Cells | 3 | 3 |
| GSE15919 | GPL1528 | (Binet *et al.* 2009) | - | MRC5 Fribroblasts | 6 | 6 |
| GSE10570 | GPL4133 | (Bhatia *et al.* 2008) | Prostate | NHP8 | 3 | 3 |

| | | | | | | |
|---|---|---|---|---|---|---|
| GSE6762 | GPL4693 | (Johung *et al.* 2007) | - | Primary Fibroblasts | 4 | 4 |
| GSE687 | GPL506 | (Hardy *et al.* 2005) | Mammary Stroma | Primary Fibroblasts | 12 | 12 |
| GSE34303 | GPL4133 | (Ren *et al.* 2013) | Bone Marrow | Stromal Cells | 20 | 20 |

**Figure S1 – GO enrichment analysis of cellular senescence signatures**

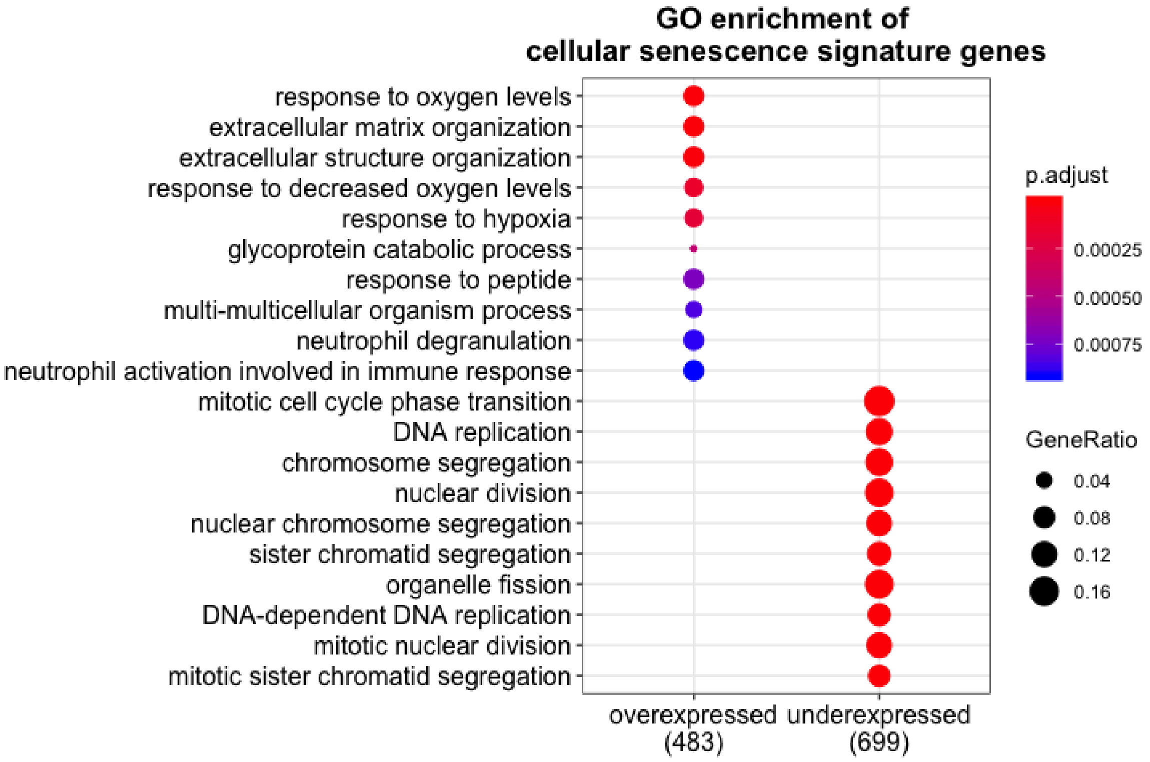

**Figure S2 – KEGG pathway enrichment analysis of cellular senescence signatures**

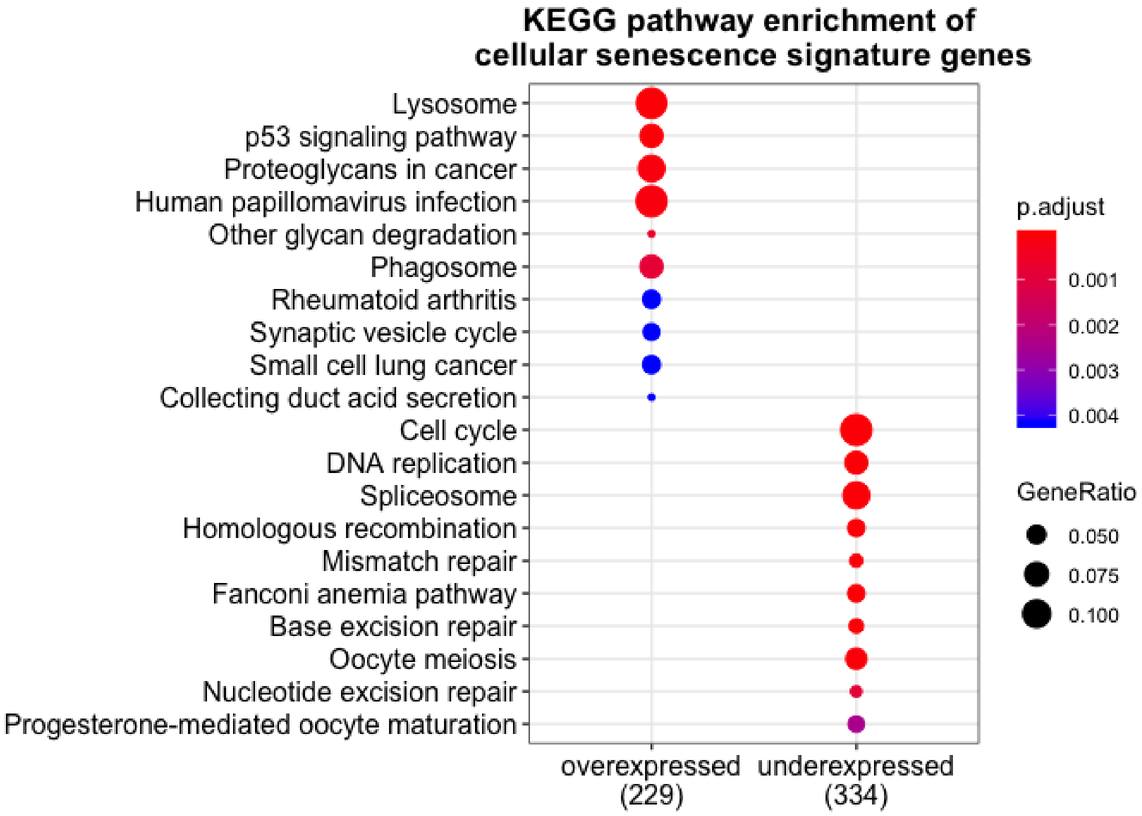

### References


Benisch P, Schilling T, Klein-Hitpass L, Frey SP, Seefried L, Raaijmakers N, . . . Jakob F (2012). The transcriptional profile of mesenchymal stem cell populations in primary osteoporosis is distinct and shows overexpression of osteogenic inhibitors. *PLoS One*. **7**, e45142.

Bhatia B, Jiang M, Suraneni M, Patrawala L, Badeaux M, Schneider-Broussard R, . . . Tang DG (2008). Critical and distinct roles of p16 and telomerase in regulating the proliferative life span of normal human prostate epithelial progenitor cells. *J Biol Chem*. **283**, 27957-27972.

Binet R, Ythier D, Robles AI, Collado M, Larrieu D, Fonti C, . . . Pedeux R (2009). WNT16B is a new marker of cellular senescence that regulates p53 activity and the phosphoinositide 3-kinase/AKT pathway. *Cancer Res*. **69**, 9183-9191.

Cancer Genome Atlas N (2012a). Comprehensive molecular characterization of human colon and rectal cancer. *Nature*. **487**, 330-337.

Cancer Genome Atlas N (2012b). Comprehensive molecular portraits of human breast tumours. *Nature*. **490**, 61-70.

Cancer Genome Atlas Research N (2014a). Comprehensive molecular characterization of gastric adenocarcinoma. *Nature*. **513**, 202-209.

Cancer Genome Atlas Research N (2014b). Comprehensive molecular profiling of lung adenocarcinoma. *Nature*. **511**, 543-550.

Cancer Genome Atlas Research N (2014c). Integrated genomic characterization of papillary thyroid carcinoma. *Cell*. **159**, 676-690.

Cancer Genome Atlas Research N (2015). The Molecular Taxonomy of Primary Prostate Cancer. *Cell*. **163**, 1011-1025.

Cancer Genome Atlas Research N, Analysis Working Group: Asan U, Agency BCC, Brigham, Women's H, Broad I, . . . Project Team: National Institutes of H (2017). Integrated genomic characterization of oesophageal carcinoma. *Nature*. **541**, 169-175.

Cancer Genome Atlas Research N, Kandoth C, Schultz N, Cherniack AD, Akbani R, Liu Y, . . . Levine DA (2013). Integrated genomic characterization of endometrial carcinoma. *Nature*. **497**, 67-73.

Cancer Genome Atlas Research Network. Electronic address wbe, Cancer Genome Atlas Research N (2017). Comprehensive and Integrative Genomic Characterization of Hepatocellular Carcinoma. *Cell*. **169**, 1327-1341 e1323.

Cao K, Blair CD, Faddah DA, Kieckhaefer JE, Olive M, Erdos MR, . . . Collins FS (2011). Progerin and telomere dysfunction collaborate to trigger cellular senescence in normal human fibroblasts. *J Clin Invest*. **121**, 2833-2844.

Consortium GT (2015). Human genomics. The Genotype-Tissue Expression (GTEx) pilot analysis: multitissue gene regulation in humans. *Science*. **348**, 648-660.

de Magalhaes JP, Curado J, Church GM (2009). Meta-analysis of age-related gene expression profiles identifies common signatures of aging. *Bioinformatics*. **25**, 875-881.

Durinck S, Spellman PT, Birney E, Huber W (2009). Mapping identifiers for the integration of genomic datasets with the R/Bioconductor package biomaRt. *Nat Protoc*. **4**, 1184-1191.

Ferreira PG, Munoz-Aguirre M, Reverter F, Sa Godinho CP, Sousa A, Amadoz A, . . . Guigo R (2018). The effects of death and post-mortem cold ischemia on human tissue transcriptomes. *Nat Commun*. **9**, 490.

Frobel J, Hemeda H, Lenz M, Abagnale G, Joussen S, Denecke B, . . . Wagner W (2014). Epigenetic rejuvenation of mesenchymal stromal cells derived from induced pluripotent stem cells. *Stem Cell Reports*. **3**, 414-422.

Gruber HE, Hoelscher GL, Ingram JA, Zinchenko N, Hanley EN, Jr. (2010). Senescent vs. non-senescent cells in the human annulus in vivo: cell harvest with laser capture microdissection and gene expression studies with microarray analysis. *BMC Biotechnol*. **10**, 5.

Guerrero A, Iglesias C, Raguz S, Floridia E, Gil J, Pombo CM, Zalvide J (2015). The cerebral cavernous malformation 3 gene is necessary for senescence induction. *Aging Cell*. **14**, 274-283.

Hardy K, Mansfield L, Mackay A, Benvenuti S, Ismail S, Arora P, . . . Jat PS (2005). Transcriptional networks and cellular senescence in human mammary fibroblasts. *Mol Biol Cell*. **16**, 943-953.

Hernandez-Segura A, de Jong TV, Melov S, Guryev V, Campisi J, Demaria M (2017). Unmasking Transcriptional Heterogeneity in Senescent Cells. *Curr Biol*. **27**, 2652-2660 e2654.

Imai Y, Takahashi A, Hanyu A, Hori S, Sato S, Naka K, . . . Hara E (2014). Crosstalk between the Rb pathway and AKT signaling forms a quiescence-senescence switch. *Cell Rep*. **7**, 194-207.

Johung K, Goodwin EC, DiMaio D (2007). Human papillomavirus E7 repression in cervical carcinoma cells initiates a transcriptional cascade driven by the retinoblastoma family, resulting in senescence. *J Virol*. **81**, 2102-2116.

Jong HL, Mustafa MR, Vanhoutte PM, AbuBakar S, Wong PF (2013). MicroRNA 299-3p modulates replicative senescence in endothelial cells. *Physiol Genomics*. **45**, 256-267.

Kim YM, Byun HO, Jee BA, Cho H, Seo YH, Kim YS, . . . Yoon G (2013). Implications of time-series gene expression profiles of replicative senescence. *Aging Cell*. **12**, 622-634.

Medeiros Tavares Marques JC, Cornelio DA, Nogueira Silbiger V, Ducati Luchessi A, de Souza S, Batistuzzo de Medeiros SR (2017). Identification of new genes associated to senescent and tumorigenic phenotypes in mesenchymal stem cells. *Sci Rep*. **7**, 17837.

Mudunuri U, Che A, Yi M, Stephens RM (2009). bioDBnet: the biological database network. *Bioinformatics*. **25**, 555-556.

Pazolli E, Luo X, Brehm S, Carbery K, Chung JJ, Prior JL, . . . Stewart SA (2009). Senescent stromal-derived osteopontin promotes preneoplastic cell growth. *Cancer Res*. **69**, 1230-1239.

Ren J, Stroncek DF, Zhao Y, Jin P, Castiello L, Civini S, . . . Sabatino M (2013). Intra-subject variability in human bone marrow stromal cell (BMSC) replicative senescence: molecular changes associated with BMSC senescence. *Stem Cell Res*. **11**, 1060-1073.

Schwarze SR, DePrimo SE, Grabert LM, Fu VX, Brooks JD, Jarrard DF (2002). Novel pathways associated with bypassing cellular senescence in human prostate epithelial cells. *J Biol Chem*. **277**, 14877-14883.

Shah PP, Donahue G, Otte GL, Capell BC, Nelson DM, Cao K, . . . Berger SL (2013). Lamin B1 depletion in senescent cells triggers large-scale changes in gene expression and the chromatin landscape. *Genes Dev*. **27**, 1787-1799.

Yang J, Huang T, Petralia F, Long Q, Zhang B, Argmann C, . . . Tu Z (2015). Synchronized age-related gene expression changes across multiple tissues in human and the link to complex diseases. *Sci Rep*. **5**, 15145.

Yu G, Wang LG, Han Y, He QY (2012). clusterProfiler: an R package for comparing biological themes among gene clusters. *OMICS*. **16**, 284-287.

Zhang H, Pan KH,  Cohen SN (2003). Senescence-specific gene expression fingerprints reveal cell-type-dependent physical clustering of up-regulated chromosomal loci. *Proc Natl Acad Sci U S A*. **100**, 3251-3256.
